# Dopamine firing plays a double role in coding reward prediction errors and signaling motivation in a working memory task

**DOI:** 10.1101/2020.05.01.071977

**Authors:** Stefania Sarno, Manuel Beirán, Joan Falcó-Roget, Gabriel Diaz-deLeon, Román Rossi-Pool, Ranulfo Romo, Néstor Parga

## Abstract

Little is known about how dopamine (DA) neuron firing rates behave in cognitively demanding decision-making tasks. Here we investigated midbrain DA activity in monkeys performing a discrimination task in which the animal had to use working memory (WM) to report which of two sequentially applied vibrotactile stimuli had the higher frequency. We found that perception was altered by an internal bias, likely generated by deterioration of the representation of the first frequency during the WM period. This bias greatly controlled the DA phasic response during the two stimulation periods, confirming that DA reward prediction errors reflected subjective stimulus perception. Contrastingly, tonic dopamine activity during WM was not affected by the bias and did not encode the stored frequency. More interestingly, both WM activity and phasic responses before the second stimulus negatively correlated with reaction times of the animal after the trial start cue and thus represented motivated behavior on a trial-by-trial basis. During WM, this motivation signal underwent a ramp-like increase. At the same time, motivation reduced noise in perception and, by decreasing the effect of the bias, improved performance, especially in difficult trials. Overall, our results show that DA activity was simultaneously involved in reward prediction, motivation and WM. Also, the ramping activity during the WM period suggests a possible DA role in stabilizing sustained cortical activity, hypothetically by increasing the gain communicated to prefrontal neurons in a motivation-dependent way.

## INTRODUCTION

Recent studies have shown that stimulation uncertainty affects the phasic responses of midbrain dopamine (DA) neurons to relevant cues during perceptual decision-making tasks. It was observed that those responses were influenced by cortical decision-making processes such as stimulus detection (1) and evidence accumulation (2), in a manner consistent with the reward prediction error (RPE) hypothesis (3, 4). Importantly, under uncertain conditions phasic DA responses exhibit beliefs about the state of the environment (3–9) and are affected by the level of uncertainty that the animal has on its choice (1, 3, 4). Although these studies represent substantial progress towards understanding how DA neurons behave under uncertain conditions, little is known about phasic and tonic (fast and slow fluctuations) DA signaling in cognitively more demanding decision-making tasks involving working memory (WM) or internal biases.

The vibrotactile frequency discrimination task (10, 11) facilitates deeper investigation into these issues. In this task, monkeys discriminated between two randomly selected vibrotactile stimulus frequencies, which were individually paired for each trial, and separated by 3 seconds (the delay, Fig. 1A). Importantly, the monkeys had to maintain a trace of the first stimulus (f_1_) during the delay period for comparison against the second stimulus (f_2_) to reach a decision: f_1_ is greater than f_2_ (f_1_>f_2_), or vice versa (f_1_<f_2_).

**Figure 1.**
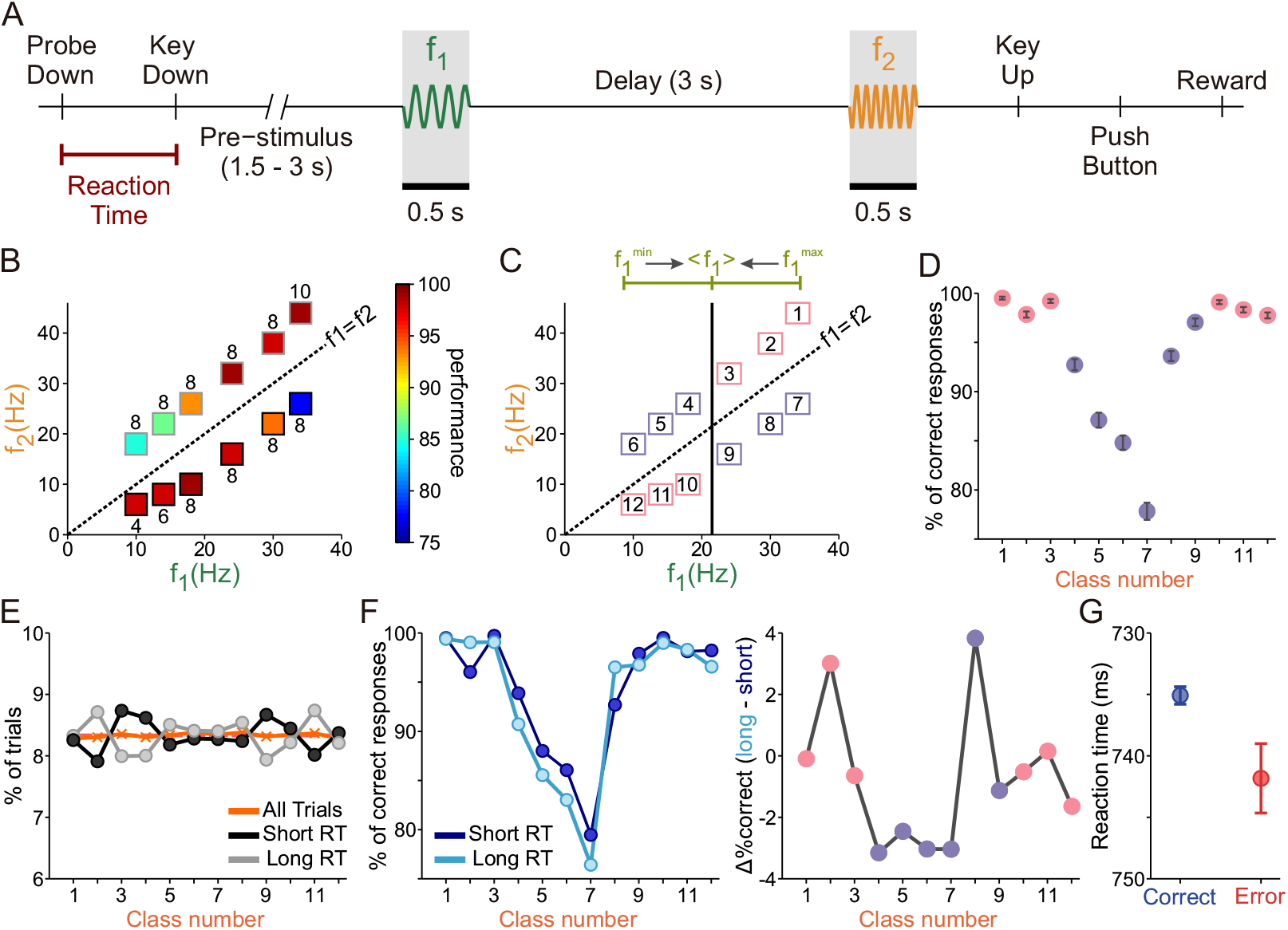
Discrimination task and contraction bias. (A) Schematic of the task design. The trial begins with the lowering of the mechanical stimulation probe (probe down, PD). The monkey reacted by placing its free hand on an immovable key (key down, KD). After a variable period (1.5-3s) the probe oscillated for 0.5s at the base (f) frequency. A 3-second delay after the first stimulus constitutes the working memory period (WM). The delay is followed by the presentation of the second stimulus (f_2_, also the “comparison period”). The offset of f_2_ signals the monkey to report its decision, and the time between offset and the monkey releasing the key (key up, KU) is known as the reaction time (RT, underlined in red). The monkey reports his decision with one of the two push buttons (PB) to indicate whether the comparison frequency was higher or lower than the base frequency. After that, the animal was rewarded for correct discriminations, or received a few seconds of delay for incorrect discriminations. (B) Stimulus set composed of frequency pairs (f_1_, f_2_) used in the task. Frequency values for f_1_ are on the x-axis, while values for f_2_ are on the y-axis. The diagonal dashed line represents stimulus equivalence (f_1_ = f_2_). The box colors indicate the percentage of correct trials, while the numbers above or below each box indicate the absolute magnitude difference (Δf) for each class. Red is for highest correct percentages; blue is for lowest. (C) The upper line (green) depicts the contraction bias effect, where the base stimulus is perceived as closer to the mean value of the first frequency (<f_1_>). The solid, vertical black line marks this center value. Numbers label classes, with the diagonal line marking stimulus equivalence (black, dashed). Upper diagonal (f_1_<f_2_) has bias benefit (pink squares) above range center, and bias obstruction (purple squares) below range center. Lower diagonal (f_1_>f_2_) has bias benefit below range center (pink), and bias obstruction above range center (purple). (D) The numbered classes (x-axis) from panel (C) by performance percentage (% of correct responses), representing the color values used in (B). Pink and purple circles represent classes with bias benefit and bias obstruction, respectively. Error bars (in black) represent standard deviations and were obtained computing the performance 1000 times, resampling with replacement from original data. (E) Percentage of trials belonging to each class number for short-RT trials (RT<P50) and long-RT trials (RT>P50; black and gray circles and line, respectively). The orange line and crosses represent the percentage of trials in each class when all trials are considered. (F) Left: Percentage of correct trials, as a function of class number, for short- and long-RT (light and dark blue circles and line, respectively). Right: Difference in the percentage of correct trials between long- and short-RT trials as a function of class number. Circles have the same color code as in panel (D). (G) Mean RT in correct (blue) and error (red) trials. Error bars represent the standard error.

Studying the time course of the DA activity while monkeys perform the discrimination task is interesting for three main reasons. First, a behavioral bias, known as the contraction bias, controlled the subjective difficulty in many instances of perceptual discrimination in both humans and rodents (12–16). Given its major role in perception we expect the contraction bias to influence the reward prediction error and the phasic DA response. Furthermore, since the bias is likely to originate from WM (12, 17, 18), it is reasonable to ask whether and how it affects the DA time course during the delay period. Second, it is known that, during the same task, frontal neurons were tuned to the identity of f_1_ during the WM delay in a time-dependent manner (11, 19, 20). Since DA neurons may be associated with frontal area responses during WM, it is a question of interest to determine whether DA neurons were tuned to the value of f_1_ or encoded reward-related information during the delay period. Finally, DA is believed to play a major role in motivated behavior (21). This role has been poorly investigated in decision making tasks. The motivational level of the animal can be indirectly measured at the beginning of each trial by considering the time it takes to react to the start cue signal and confirm its readiness to perform the task (Reaction Time, RT; see Fig. 1A). Following this logic, Satoh et al.(22) found that the phasic DA activity, besides conveying a reward prediction error signal, strongly correlated with motivation, (see also (21)). On the other hand, prior claims associated tonic DA activity with conveying motivation (23, 24). We then wondered whether motivation influences the DA transient responses and/or the working memory activity during the discrimination task.

We found that the DA activity simultaneously coded reward prediction and motivation. Phasic responses to the stimuli exhibited reward prediction errors and were shaped by the contraction bias. DA responses to the starting cue and the first stimulus correlated negatively with the RTs and thus reflected motivation on a trial-by-trial basis. WM activity conveyed a pure motivational signal and exhibited a ramping-like increase during the delay period, which was more pronounced in high-motivation trials (trials with a shorter RT), exhibiting correlations with RTs on single trials. At a behavioral level, motivation seemed to reduce noise in perception and, by decreasing the effect of the bias, improved performance, especially in difficult trials.

Overall, our study extends the well-known role of DA neurons in processing RPEs over belief states to tasks that present an internal bias. It also shows that willingness to work for rewards leads to better outcomes by enhancing precision and reducing a perceptual bias. More importantly it provides strong evidence that, in well-trained animals and in absence of any external cue, the activity of the same neurons ramps up in a motivation-dependent way during the WM delay interval. Such an increased, sustained DA activity signal may be related to a better usage of cognitive resources such as working memory.

## RESULTS

### Contraction bias and motivation shape behavioral performance

Monkeys were trained to discriminate between two vibrotactile stimuli with frequencies within the flutter range. Each stimulus pair (f_1_, f_2_; class) consisted of a large, unambiguous difference (8 Hz in most classes; Fig. 1B). In Fig. 1B we show that performance for each class was only partially predictable from the absolute value Δf = | f_1_ – f_2_ |. The contraction bias could explain this disparity, since the decision accuracy depended on the specific stimuli that were paired. This internal process shifts the perceived frequency of the base stimulus f_1_ towards the center of the stimuli frequency range (<f_1_>=21.5 Hz, Fig. 1C). As such, if f_1_ has a low stimulus value (i.e., 10 Hz), it will be perceived as larger, while a large frequency for f_1_ is perceived as smaller. Therefore, correct evaluations for classes with f_1_<f_2_ are increasingly obfuscated by the bias’s effect as classes are visited from right to left (upper diagonal, Fig. 1B-C), while classes with f_1_>f_2_ are increasingly facilitated by the bias (lower diagonal, Fig. 1B-C). To preserve this bias structure, we labeled classes with beneficial bias effects as closer to the extreme ends, and classes with hindrance from the bias as the center classes (Fig. 1C). In this class organization, decision accuracy was highest for the classes near to the two ends (bias benefit) and, when plotted as a function of class number, the resulting curve was U-shaped (Fig. 1D).

To determine whether the motivation of the animals affected their behavioral performance we analyzed the influence of RT on accuracy. First, we classified trials according to RT into two groups (short & long, Methods) and we observed that, in both conditions, the distribution of trials as a function of class number was not significantly different from the uniform distribution (see Fig. 1E; Chi-Squared test, p=0.46). This result confirmed that, at the beginning of the trial, the animal had no clue about the upcoming class and thus that the RT can be considered as a behavioral measure of motivation independent of the difficulty.

Since motivation can affect perception (25, 26) we next considered how the RT was related to accuracy. We observed that accuracy for long RT was lower than for short RT (Fisher’s exact test, p=0.036). As a function of class number (Fig. 1F, left), in both short- and long-RT trials, the accuracy decreased for classes disfavored by the bias (classes 4-9, Fig. 1C purple). Furthermore, in those same classes, the performance was significantly better for short-RT trials (Fig. 1F, right; Fisher’s exact test, p=0.003); on the contrary, differences in accuracy did not reach a significant level (Fisher’s exact test, p=0.88) in classes that were favored by the bias (classes 1-3 & 10-12, Fig. 1C pink). Finally, we compared the RT in correct and error trials (Fig. 1G) and observed that the RT was significantly shorter in correct trials (t-test, p=0.009).

All the above results indicated that the motivation had a strong effect on the performance of the animal: It strongly correlated with the trial outcome and significantly improved the accuracy in subjectively difficult trials (those corresponding to classes disfavored by the bias).

### Contraction bias affects DA responses

We recorded single-unit activity from 22 putative midbrain DA neurons while trained monkeys performed the discrimination task. They were identified based on previous electrophysiological criteria (27): regular and low tonic firing rates (mean ± SD = 4.7 ± 1.4 spikes/s), a long extracellular spike potential (2.4 ms ± 0.4 ms), positive activation to reward delivery in correct (rewarded) trials and with a pause in error (unrewarded) trials (28–30).

We found that the majority of neurons (16 out of 22, 72% of the population) showed a significant, positive modulation to at least one of the two stimuli (see Methods). We therefore analyzed how the DA neurons responded during the stimulation period. It is known that, in frontal lobe neurons, f_1_ is parametrically represented during WM and the animal’s decision is observable during the comparison period (i.e. during the presentation of f_2_; (31)). We hypothesized that if DA neurons receive feedback information from the frontal lobe, their responses to f_2_ should reflect the contraction bias’s effect as part of the decision report. Resultantly, neurons showed significant dependence on stimulus classes (one-way ANOVA, p=0.0013; Fig. 2A). The U-shaped modulation in the response distributions can be explained by a pair of arguments: first, the bias induced a subjective difficulty that affected belief states. Second, as an RPE signal, the DA response to f_2_ coded the change in reward expectation produced by the application of the comparison frequency, which is higher for classes facilitated by the bias (Fig. 2A). A reinforcement learning model (32) based on belief states supports this interpretation (Fig. S1A). Similar effects of belief states on responses to reward-predicting stimuli were observed in other experiments (2–4). However, in those works the trial difficulty was controlled by experimental design while it was determined by a perceptual bias in our task.

**Figure 2.**
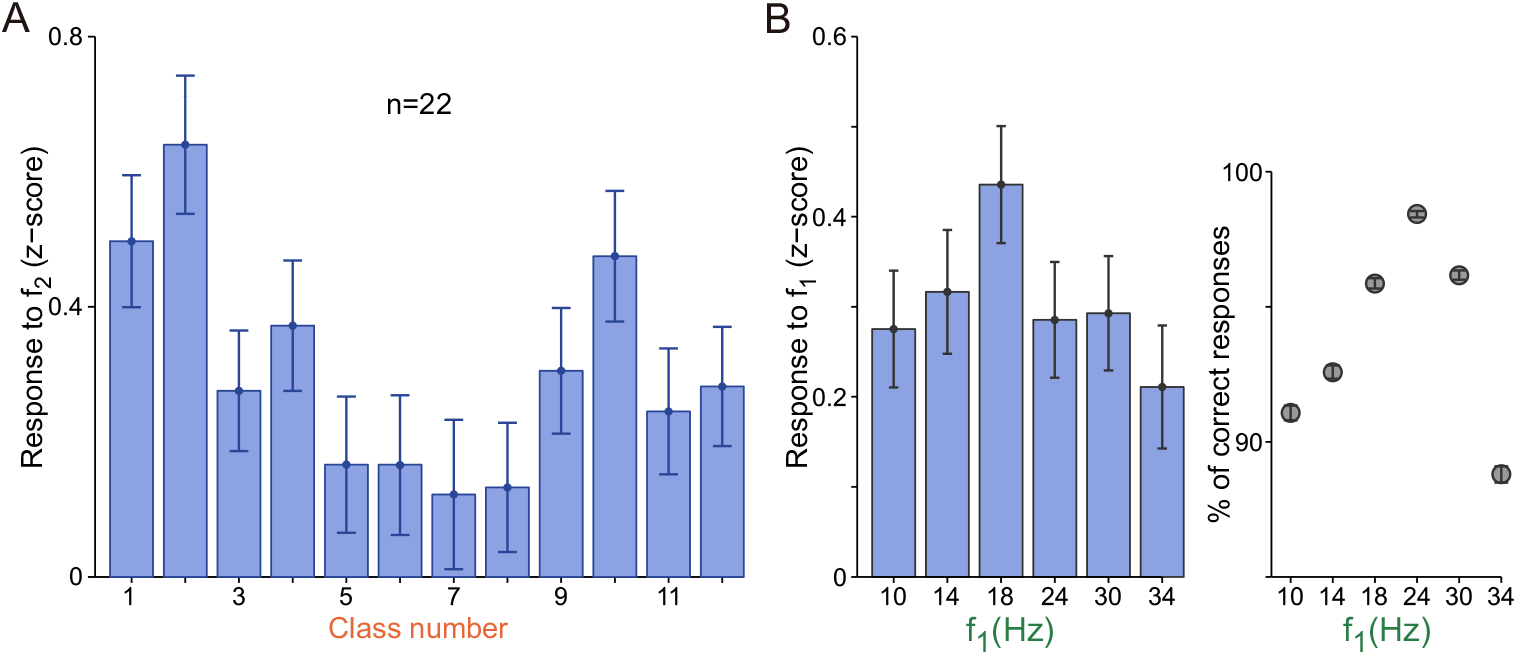
(A) Normalized population responses to f_2_, sorted by stimulus class (Methods). Error bars are ±1 SEM. (B) (Left) Normalized population response to f_1_ plotted against f_1_ frequencies. Error bars are ± 1 SEM. (Right) Percentage of correct responses for each f_1_ frequency.

Next, we wondered whether the same bias effect was observable during f_1_. By design, each f_1_ value is the same for two classes (f_1_>f_2_ or f_1_<f_2_), where one class is favored by the bias, and the other is disfavored (Fig. 1D). However, as the animal had no clue about the upcoming class during the presentation of the first stimulus, it is reasonable to study the monkeys’ performance and DA responses based on the f_1_ value only. We observed that the decision accuracy was lowest for the extremum f_1_ intensities (Fig. 2B, Right). As we would expect, the phasic DA response to f_1_ showed a similar modulation (Fig. 2B). Furthermore, DA responses for the middle values of our stimulus range (18 and 24 Hz) were significantly greater than those produced after the application of extreme values of the stimulus range (i.e. for 10 and 34 Hz; p = 0.039, two sample one-tailed t-test).

### Contraction bias determines the subjective difficulty

We reasoned that the contraction bias could influence the animal’s subjective difficulty of a class, similarly as the stimulus physical features do it (see, e.g., (33)). To analyze this issue, we used a Bayesian approach in which observation probabilities were combined with prior knowledge to obtain a posterior probability or belief about the state of the stimuli (13, 17, 34–36) (Fig. 3A). The prior probabilities of f_1_ and f_2_ were assumed to be discrete and uniform, and we supposed that animals only perceived noisy representations of the two stimuli (Fig. 3A). Observations of f_1_ and f_2_ were obtained from Gaussian distributions at the end of the delay period, with respective standard deviations σ_1_ and σ_2_ as noise parameters. Decisions were made using the belief about the state f_1_>f_2_ and the *maximum a posteriori* (MAP) decision rule. The noise parameters were adjusted to optimize the similarity between the Bayesian model and the animal’s performance (Fig. 3B, see Methods for model and parameter fitting). The best fit yielded σ_2_=3.2 Hz and σ_1_=5.50 Hz (Fig. 3B), so the noise parameter for f_1_ was greater than that for f_2_; from this, we intuited that the representation of f_1_ deteriorated over the WM period. When this happens, the Bayesian model produces a contraction bias because the inferred value of f_1_ is noisier than that of f_2_, and the inferred value of f_1_ relies more heavily on its prior distribution (13).

**Figure 3.**
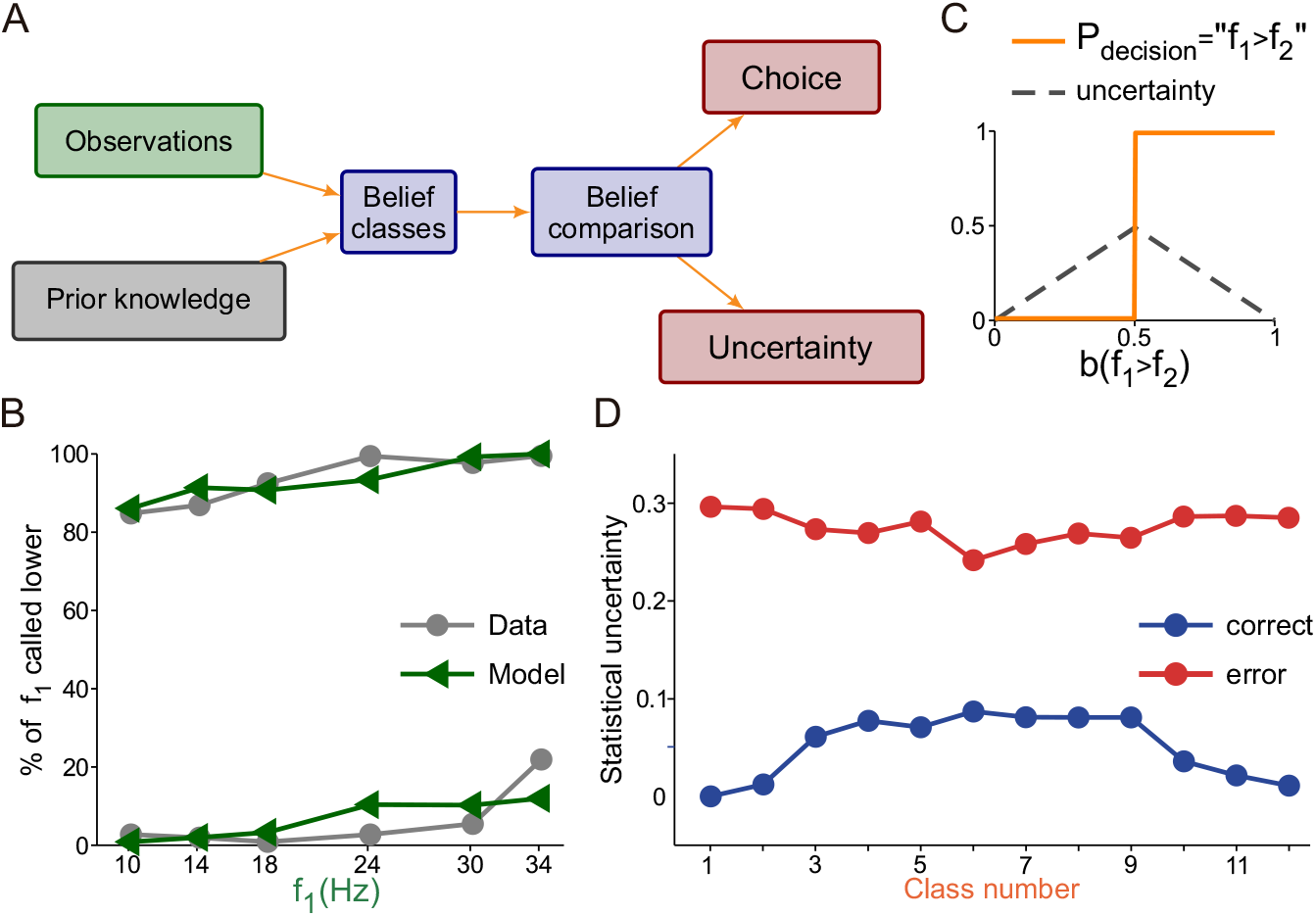
A Bayesian behavioral model and the relationship between the contraction bias and confidence. (A) Schematic of the Bayesian Behavioral Model: prior knowledge and observations are combined to obtain two comparison beliefs, b(H) (f_1_>f_2_, Higher), and b(L) (f_1_<f_2_, Lower). These beliefs are compared by magnitude in order to reach a decision. The decision can be decomposed into two components: Choice and Uncertainty. See Methods for a more accurate description of the model. (B) Model fit of the percentage of times in which the base frequency (f_1_) is called smaller. The behavioral data in grey (circles, lines), model predictions in green (triangles, lines). The line pairs for the 12 classes divided by the diagonal in Fig. 1D (f_1_<f_2_, upper pair; f_1_>f_2_, lower pair). Model best-fit parameter values were σ_1_ = 5.5 Hz and σ_2_ = 3.2 Hz. (C) Choices are made using maximum a posteriori (MAP) criterion (orange line). Uncertainty is defined as one minus the maximum of the two beliefs b(H) and b(L) (grey line). (D) Bayesian model uncertainty predictions: correct trials in blue (circles, lines) and wrong trials in red (circles, lines).

We then used the fitted Bayesian behavioral model to simulate the discrimination task and to evaluate the subjective difficulty for each trial. Subjective difficulty was defined as the decision uncertainty (33), given by the probability that the choice made by the model was not correct (Fig. 3C). The subjective difficulty associated with a class was defined as its average over many simulated trials in that class. In correct trials, class difficulty versus class number was an inverted U (Fig. 3D, blue line), increasing/decreasing as the bias was less/more favorable to making a correct choice. The two classes (11 and 12) with smaller absolute magnitude difference Δf (6Hz and 4Hz) had decreased class difficulty due to the bias, although the increased difficulty in discrimination somewhat compensates its effect. Class 1 was doubly facilitated: the bias decreased class difficulty level and it had the largest Δf (10Hz), and as such the difficulty of discrimination was less. Comparing the results between correct and error trials, the model had the opposite class subjective difficulty pattern when it erred (Fig. 3D, red line). Thus the contraction bias regulated class difficulty in a way similar to how an odor mixture modulated the choice uncertainty of rodents executing an odor categorization task (33). However, the important distinction for our results is that subjective difficulty was mainly modulated by an internally generated process, and not by the physical properties of the stimuli.

### Subjective difficulty shapes DA responses during decision formation

To study the influence of class subjective difficulty on DA activity, we considered correct trials and subdivided the stimulus set (12 classes) into two groups: a low-difficulty group (classes 1-3 & 10-12, Fig. 1C pink) and a high-difficulty group (classes 4-9, Fig. 1C purple). During presentation of f_1_, the normalized activity (z-score) showed a tendency to be higher in the low-difficulty group (Fig 4A left); however, this tendency was not found to be significant (p=0.17, two tail t-test; Methods). Importantly, the difference became significant during the presentation of f_2_ (Fig. 4A, center; p<0.001, two tail t-test; Methods). We obtained a response latency of 245ms for the time it took the subjective difficulty groups to significantly diverge in responses after f_2_ onset (AUROC, p<0.05, sliding 250 ms window, 10 ms steps; Methods). Dependence on class difficulty during decision formation is similar to the dependence on choice confidence as reported in previous works (3, 4). Interestingly, the activity after delivery of reward was invariant to the subjective difficulty level, having the same peak response for both class groups (Fig. 4A, right).

**Figure 4.**
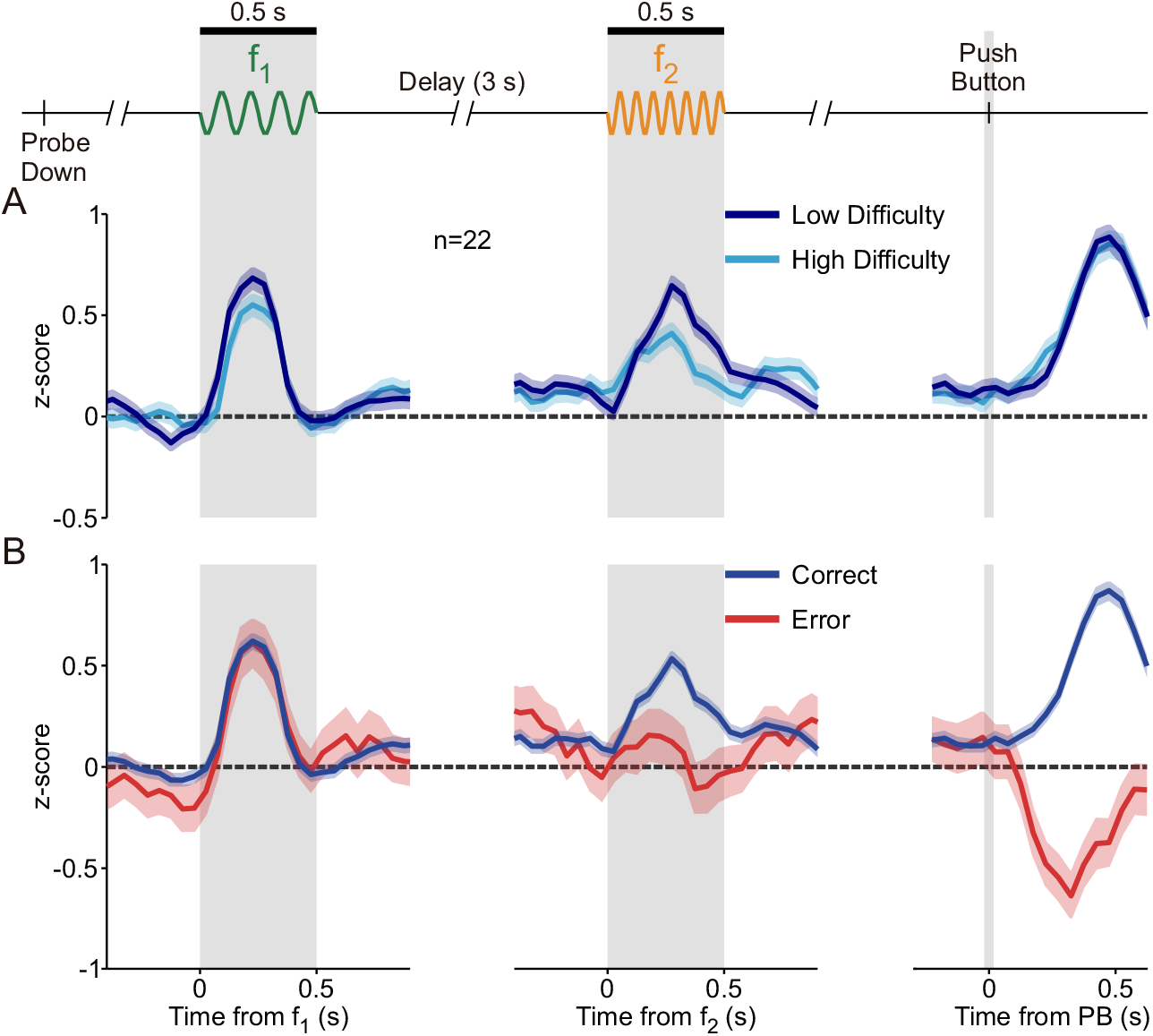
The phasic activation of DA neurons to the base stimulus, comparison stimulus, and reward delivery. Normalized DA population responses to different periods of the task (shadows ±1 SEM): (Left) Responses to f_1_ (grey shaded region), aligned to f_1_ onset. (Center) Responses to f_2_ (grey shaded region), aligned to f_2_ onset. (Right) Responses to reward delivery, after the decision report (slender grey region), aligned to PB. (A) Responses for confidence groups: high (navy blue) and low (light blue) confidence. (B) Responses for trial outcome groups: correct (blue) and error (red) trials.

The Bayesian model predicted higher subjective difficulty for error trials (Fig. 3D), so we studied the phasic activity sorting trials by decision outcome (correct vs. incorrect). Responses to f_1_ did not vary based on outcome (Fig. 4B, left), but the responses did vary during the comparison stimulus. The temporal profiles for each outcome differentiated with an analogous latency value (~205ms) to the class difficulty groups (Fig. 4B, center). This time lag in DA signals is comparable with those found in the secondary somatosensory cortex (37) and frontal areas during the same task (31). Similarities in latency between frontal lobe and DA neurons were also found during a tactile detection task (38).

At the point of reward delivery, the phasic DA activity increased in correct trials, but decreased for error trials (Fig. 4B, right). This temporal pattern is consistent with that predicted by the RPE hypothesis. The latency of divergence of these two response profiles (AUROC, p<0.05, Methods) was shorter than those after f_2_ onset (~130ms). Further, we asked whether a reinforcement learning model based on belief states could emulate the dopamine responses. The model reproduced the phasic responses observed in Fig. 4 well (Fig. S1B-C).

### Dopamine ramps up during the delay WM period without coding f_1_

Because during the WM period neurons in frontal areas exhibit a temporally modulated sustained activity, tuned parametrically to f_1_ (11, 19, 31), we investigated how DA neurons behave during the same period. Is their activity modulated and tuned to f_1_? Fig. 5A (left) shows the normalized activity (Methods) of an example neuron with significantly increased activity through the entire WM period. When we sorted the firing rates by f_1_ values, we saw a clear ramping in activity across the delay period in all 6 curves (Fig. 5A, center). However, the temporal profiles are not modulated by the f_1_ values. To further verify the absence of tuning, we averaged the entire WM activity for each f_1_ value (Fig. 5A, right), and found that this neuron did not exhibit significant differences across f_1_ (one-way ANOVA, p = 0.76). Fig. 5B (left) shows another example neuron. Its activity exhibited absolutely no temporal modulation during WM, remaining closer to its baseline value (Fig. 5B, left). The absence of ramping persisted when we expanded the observations to differentiate between responses to each f_1_ stimulus, (Fig. 5B, center). Further, neither the temporal profile nor the averaged activity during WM (Fig. 5B, right) were modulated based on f_1_ (one-way ANOVA, p = 0.64).

**Figure 5.**
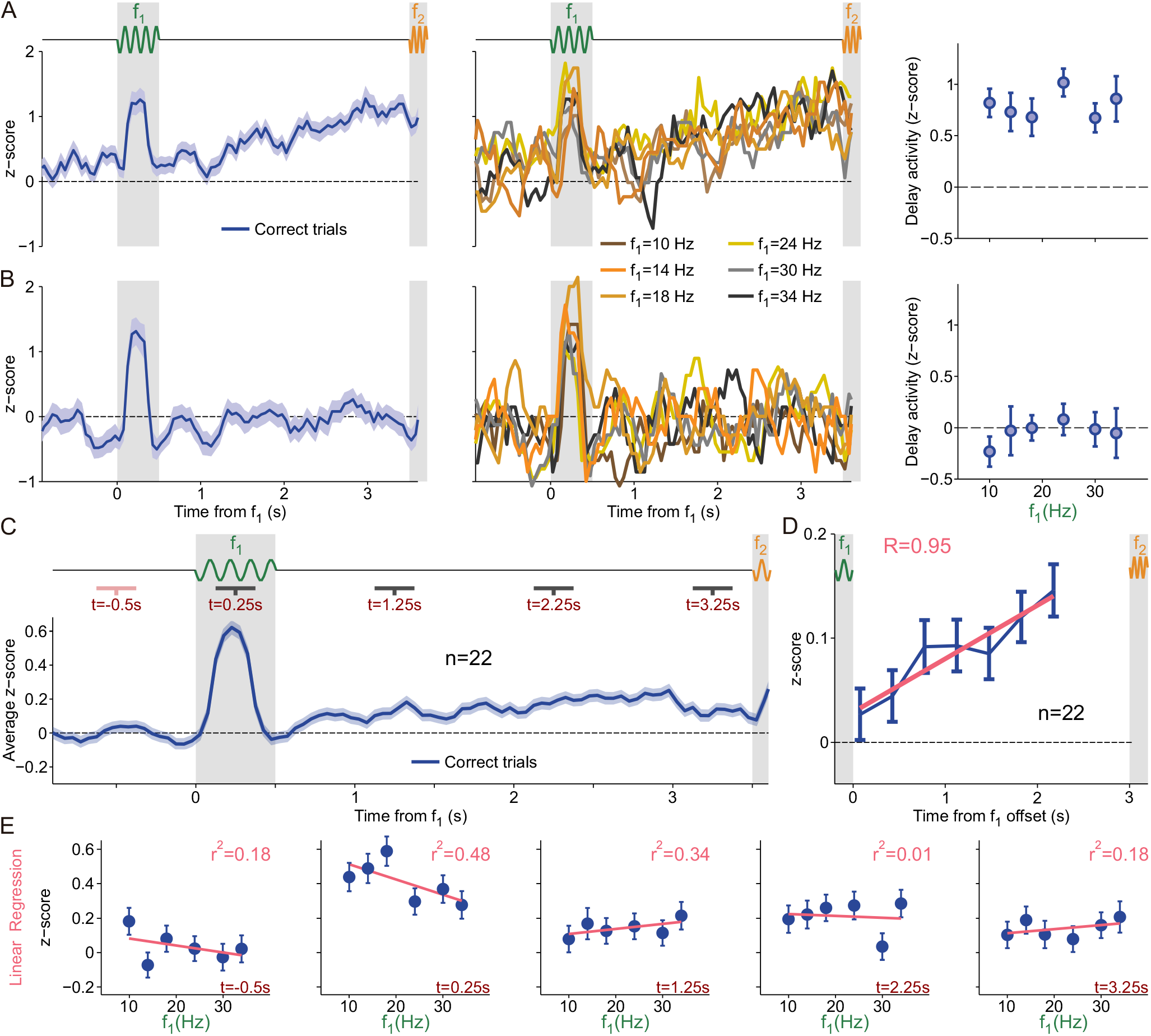
DA Activity during the delay period. (A-B) Normalized activity as a function of time for 2 example neurons, trials aligned to f_1_ onset. Activity calculated in windows of 250ms with steps of 50ms (Methods). Neither example shows coding activity during WM. (Left) Activity from all correct trials was averaged (blue line, shadows ±1 SEM). Thick grey shaded area for f_1_, thin grey shaded region for f_2_. (Center) Same marked grey regions. Each curve indicates activity for one f_1_ stimulus value. Curves with color closest to yellow for intermediate values, darkest curves for the extreme values. (Right) Average activity during the WM (delay) period as a function of f_1_ values (error bars ±1 SEM). (A) Exemplary neuron with temporal dynamics during WM. (B) Single neuron without temporal dynamic. (C) Normalized activity as a function of time between 1 second before f_1_ onset through the end of WM (blue line, shadows ±1 SEM). Trials aligned to f_1_ onset. Grey regions mark the same periods as for (A-B), activity calculated in the same manner. Dark grey lines under task indicate analyzed periods after f_1_ onset, pink line under task indicates the basal period before f_1_ onset. Pink periods compared against grey periods show significant modulations in each time segment (Wilcoxon signed-rank test; p<0.001, p=0.012, p<0.001 and p<0.01). (D) Mean population activity of DA neurons during the delay period, with correction for basal activity. Grey shadowed regions demonstrate stimulus presentation periods. The blue points for corrected firing rate in non-overlapping 350ms periods (±1 SEM). Pink line displays linear regression across 8 discrete periods, where R is the determination coefficient. (E) Linear regressions (pink line) calculated in each of the 5 indicated regions from panel (C). Regressions over average activity for each f_1_ stimulus (error bars for ±1 SEM). First graphic for the basal period before f_1_ onset, proceeding in increasing chronological order from left to right. Times in the bottom left corner (dark red), r^2^ values are the determination coefficients and show the accuracy of each linear fit to the data (upper right corner, light red). No linear regressions were found to have significant monotonic coding.

At the population level, we found that neurons responded to f_1_ by increasing their firing rate in a way that was not tuned to the value of the frequency (one-tailed Wilcoxon signed-rank test; p<0.001; Fig. 5C). Furthermore, dividing the WM period into 3 segments (Fig. 5C, black horizontal bars indicated in the delay period), we observed positively and significantly modulated activity in all of them when compared to a segment before the presentation of f_1_ (Fig. 5C, pink bar; one-tailed Wilcoxon signed-rank test; p=0.012, p<0.001 and p<0.01 respectively). The average population activity exhibited a ramping increase during the WM period (Fig. 5D).

Due to the fact that frontal neurons can be monotonically tuned to the identity of f_1_ during WM, we sought to confirm that this coding did not exist within the DA population in any time interval. A linear regression in five discrete intervals (pink and black bars in Fig. 5C, top) showed no significant linear trend (Fig. 5E). Moreover, we extended our analysis to one second before f_1_ onset through the end of the WM (a sliding 250ms window with 10ms steps). In these 4.5 seconds, we performed linear and sigmoidal regressions in each window and found no significant results for either linear or sigmoidal regression, in any window (slope different from zero, p<0.01, and fit with Q>0.05; Methods). Notably, the absence of monotonic dependence was held even during the last portion of the delay interval. This contrasts with the f_1_ information recovery observed in frontal neurons (20), a result also found in other tactile tasks (39). Pure temporal ramping signals during WM were also identified in frontal areas in this and another tactile tasks (19, 40).

To exclude the existence of more general dependencies on f_1_ during WM (e.g., similar to those in Fig. 2) we searched for temporal windows with significant ANOVA results (p<0.05; Methods). We focused on the same 4.5 seconds and found that in several moments during the presentation of f_1_ and throughout the delay period the activity depended on f_1_; however, this dependence was intermittent without exhibiting a clear or consistent pattern (Fig. S2A-B). This led us to seek temporal consistency, so we divided the entire time range into 9 discrete subintervals of 500 ms and calculated the fraction of significant sliding windows in each (Methods). We found a consistent dependence on f_1_ only during the presentation of the stimulus, where the z-scores depended on f_1_ with a profile similar to the one shown in Fig. 2B (left) (see also Fig. S2B).

After studying f_1_ in isolation, we repeated a similar analysis to identify dependencies on the stimulus class. We implemented the ANOVA test (see Fig. S2C) and searched for those windows in which p<0.05. We then calculated the temporal consistency in discrete subintervals of 500ms, repeating the same procedure as for f_1_. We found significant class-dependent activity only during f_2_ stimulation, with a pattern similar to that in Fig. 2A (see Fig. S2D and Methods).

### Phasic DA activity reflected motivation on a trial-by-trial basis only before the decision formation

To understand the influence of motivation on a trial-by-trial basis we first classified trials according to RT into two groups (short and long, Methods) and analyzed the time courses of DA activity using this classification (Fig. 6A). We noticed that the phasic responses to the PD and to f_1_ were higher for short-RT trials (Fig. 6A, left and middle). Contrastingly, both groups reached a common peak response to the second stimulus (Fig. 6A, middle) and to the reward delivery (Fig. 6A, right).

**Figure 6.**
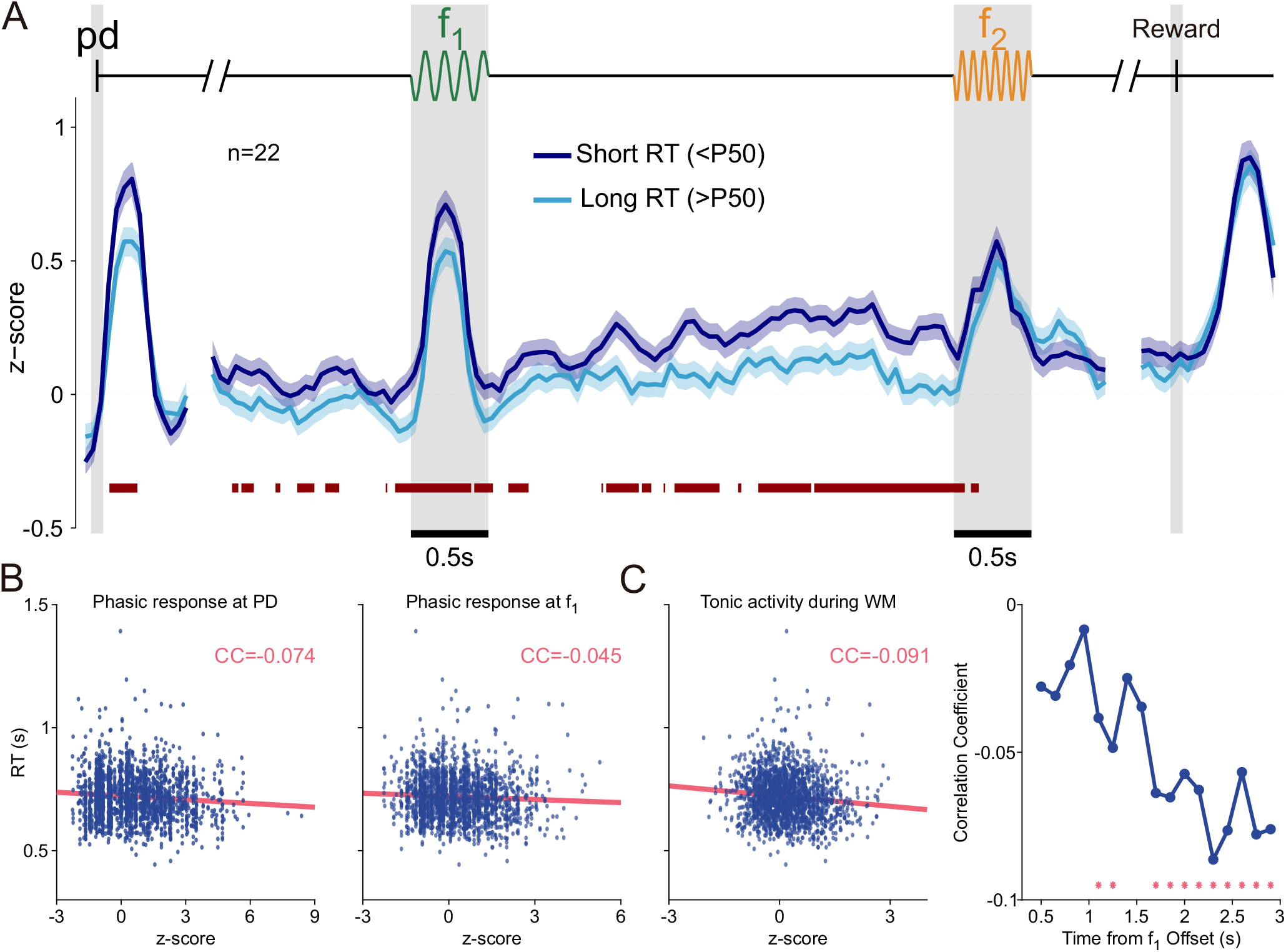
DA activity in short- and long-RT trials. (A) Normalized population activity as a function of time for correct trials, divided by reaction time duration: short-RT trials (RT<P50 (<50%), dark blue line, shadows ±1 SEM) and long-RT trials (RT>P50 (>50%), light blue line, shadows ±1 SEM). Red colored bar below curves marks significant differences between RT groups (AUROC, p<0.05; Methods). (B) Trial-to-trial correlation of the magnitude of the DA response to the PD (left) and to f_1_ (right) with the RTs. (C) (Left) Trial-to-trial correlation between the DA activity during the WM and the RTs. (Right) Evolution of the correlation coefficient between the DA activity and the RTs through the WM period. Red stars mark windows with significant correlation coefficients. Note that correlations were consistently negative and became significant towards the end of the delay WM period.

This analysis suggests that, prior to the presentation of f_2_ and to the decision formation, the phasic DA responses to the PD and f_1_ coded motivation (Fig. 6A). To confirm this result, we computed the correlation coefficient between the RTs and the phasic DA responses. We obtained a significant negative correlation between the DA responses to the PD and the RTs (r=-0.07, p<0.01 with permutation test; Fig. 6B left and Methods). A similar result was found when we analyzed the correlations between the RTs and the DA activity after the first stimulus (r=-0.05, p=0.021 with permutation test; Fig 6B right). Contrastingly, the correlations were not significant when we considered the responses to the second stimulus and reward delivery. This suggests that during and after the decision formation, the DA phasic activity ceased to code motivation and only reflected a RPE signal.

### Tonic DA activity increasingly coded motivation throughout the WM period

We finally analyzed how the DA activity varied with the RT during the WM period. We found that for short-RT trials the tonic DA signal (z-score) was consistently greater than for long-RT trials. The red bar in Fig. 6A indicates periods of significant differences between these two signals (AUROC, p < 0.05). When we averaged the activity for the RT groups across the whole WM period, the activity of short-RT trials was significantly higher ([mean ± SEM]_short_=0.25±0.03; [mean ± SEM]_long_=0.11±0.03; p<0.001, two-sample one-tailed t-test). Furthermore, during WM, the trial-to-trial correlations between the DA activity and the RT were significantly negative (r=-0.10, p=<0.001 with permutation test; Fig 6C left and Methods). More interestingly we noticed that the DA activity in short- and long-RT trials became increasingly different as time elapsed during the delay period, showing a higher value in short-RT trials (Fig 6A). To assess the significance of this effect we divided the delay period in nonoverlapping bins and computed the correlations between the DA activity and the RT in each of them (see Methods). We found that the correlation coefficient consistently decreased, becoming significantly negative during the last portion of the delay (Fig 6C right, see Methods). To summarize: the influence of motivation on DA activity was more pronounced at the end of the WM period, while tonic DA simultaneously ramped up.

### Motivation improved perceptual precision

We have previously noticed that the performance was better in short-RT trials, especially in classes negatively affected by the bias (Fig.1F). We thus explored how this performance enhancement arose in the Bayesian framework by fitting the model parameters to both trial groups independently (see Fig.S3). We observed that the noise parameter σ_1_ (emulating the uncertainty in the representation of the memorized frequency f_1_) was smaller in the short-RT condition (p<0.001, T-test). A smaller σ_1_ decreases the contribution of the prior distribution to the posterior of f_1_, resulting in a weaker contraction bias. Thus, in short-RT trials, the performance improved in classes disfavored by the bias. Instead, in classes favored by the bias, the reduction of the uncertainty parameter had almost no effect, because the discrimination is already very easy.

## DISCUSSION

We sought to determine the activity of DA neurons recorded while monkeys performed a cognitively demanding discrimination task. We found that phasic DA responses before the comparison period coded RPE and motivation, although the dopamine activity during WM only coded motivation. We also observed that a subset of DA neurons exhibited a motivation-dependent ramping increase during the delay.

The contraction bias affected the subjective task difficulty. At the end of the delay period, observations of the first frequency have deteriorated, increasing the relevance of prior knowledge, and thus modulating performance (Fig. 1D) and DA responses (Fig. 2A). The effect of the bias was somewhat countered by motivation. A larger motivation increased the perceptual precision of the observations of the first frequency, producing a weaker contraction bias (because the prior is less relevant), reducing the subjective difficulty and boosting performance in short-RT trials (Fig. 1F). Recently, Mikhael et al. (41) proposed that tonic DA controls how worthwhile it is to pay a cognitive cost that improves Bayesian inference by increasing perceptual precision, ultimately leading to better performances. Motivation also influenced DA activity: The phasic responses to the start cue (the PD event) and the first vibrotactile stimulus depended on the RT, exhibiting a larger value for faster responses. Importantly, DA responses to those task events and the delay period activity were negatively correlated with the RT, suggesting that DA drives motivated behavior on single trials, as it has been advocated in other quite different tasks (21, 22).

An interesting result was that DA neurons exhibited RT-dependent ramping activity during WM. This DA signal conveys motivational information, which could be playing relevant roles in cortical cognitive processes. Previous recordings in midbrain DA neurons during monkeys’ performance of WM tasks did not observe sustained activity during the delay (42–44). This discrepancy can be attributed to differences in WM and attention requirements, as well as to task difficulty or delay duration. On the other hand, to our knowledge, the ramping WM activity we observed has not been reported before. In the striatum, however, a ramping increase in the release of DA has been described (21, 23, 45–47); this signal purportedly reflects an approaching decision report (23, 45), reward proximity (46, 48), possibly aiding in action invigoration as well (49). Ramping in the spiking activity of DA neurons was observed in some experiments (48, 50) but not in others (45). Moreover several theoretical explanations have been proposed for generating ramping signals (51–54). Ramping has been associated with lack of training (55) or the presence of external cues reducing uncertainty (54). Here we observed a sustained ramping of DA activity in well-trained animals and in absence of any external feedback able to reduce temporal or sensory uncertainty. The ramping DA signal appeared when an active maintenance of task-relevant information was required, and as such, we attribute its existence to the usage of WM.

Although in our experiment we were not able to address the implications of the delay period DA signal, we propose that this activity might be related to the cognitive effort needed to upload and maintain a percept in WM. Recent advances in the study of motivation and cognitive control (56) indicate possible directions to investigate this issue further in future work. Cognitive control is effortful (57, 58), involving a cost-benefit decision-making process. A task that involves WM requires an evaluative choice: either engaging in the task, or choose a less demanding option (e.g., guessing), even if it leads to a smaller reward (49, 59). An intriguing hypothesis relates motivation with cognitive control modulated by dopaminergic signals to prefrontal cortex and striatal neurons (56, 60, 61). In this scenario DA is proposed as a persistent-activity stabilizer, increasing the gain in target neurons, and in turn producing an enhanced signal-to-noise ratio and promoting WM stability (62–64). Summarily, our results underline the essential role of midbrain DA neurons in learning, motivation and working memory under perceptual uncertainty. In our frequency discrimination task, phasic DA responses were affected by the contraction bias and motivation, while working memory activity could instead be compatible with motivational behavior. The dependence of DA activity on the contraction bias suggests that internally generated biases can play a role in learning. Two-interval forced-choice tasks, such as discriminating between two temporal intervals exhibit the contraction bias (14); however, DA was not investigated in these cases. Future study within these task paradigms can further our understanding of DA activity in motivation, cognitive control and working memory. Altogether, our results point to an intricate relationship between DA, perception and WM, as they are all modulated by the animal’s intrinsic motivation.

## EXPERIMENTAL PROCEDURES

All protocols were approved by the Institutional Animal Care and Use Committee of the Instituto de Fisiología Celular, Universidad Nacional Autónoma de México.

## ACKNOWLEDGMENTS

This work was supported by Grants PGC2018-101992-B-I00 from the Spanish Ministry of Science, Innovation and Universities (to S.S., M.B. and N.P.) and by the Dirección de Asuntos del Personal Académico de la Universidad Nacional Autónoma de México, Consejo Nacional de Ciencia y Tecnología PAPIIT-IN210819 (to R.R.-P.).

## AUTHOR CONTRIBUTIONS

R.R., G.G.-d.L. and R.R.-P. performed experiments. S.S., M.B., J.F.R. and N.P. analyzed the data and designed the models. S.S. and M.B. wrote the codes. N.P., S.S, J.F.R, R.R-P and R.R. wrote the paper helped by the other authors.

## METHODS

### Discrimination Task

This study was performed on two male monkeys, Macaca mulatta, 5–7 kg. The sensory discrimination task used here has been described previously (11, 20, 37). The schematic representation of this task is depicted in Fig. 1A. The monkey sat on a primate chair with its head fixed. The right hand was restricted through a half-cast and kept in palm-up position. The left hand operated an immovable key (elbow at ~90°) and two push buttons in front of the animal, 25 cm away from the shoulder at eye level. The centers of the switches were located 7 and 10.5 cm to the left of the midsagittal plane. In all trials, the monkey first placed the left hand on the key, and later projected to one of the two switches. Trials began when the mechanical stimulator is lowered, indenting the fingertip of one digit of the restrained hand (Probe Down, PD). The monkey places its free hand on an immovable key (Key Down, KD). The time lag between PD and KD constitutes the reaction time (RT, Fig. 1A). After the KD, a variable delay of 1.5-3s is presented to avoid anticipatory activity before the arrival of the stimulus, followed by the first stimulus (f_1_), lasting 0.5s. The second stimulus (f_2_) is presented after a 3s delay, also lasting 0.5s. The offset of f_2_ signals the monkey to release the key (Key Up, KU), and report its decision by pressing one of two push buttons (PB) with the left hand (lateral push button for f_2_>f_1_, medial push button for f_2_<f_1_). Immediately after the decision report, correct discriminations were rewarded with a few drops of liquid, while incorrect discriminations received a few seconds of delay before the beginning of the next trial. Stimuli were delivered to the skin of the distal segment of one digit of the restrained right hand, via a computer-controlled stimulator (BME Systems; 2 mm round tip). Initial probe indentation was 500 μm. Vibrotactile stimuli were mechanical sinusoids pulses lasting 20ms each. Stimulation amplitudes were adjusted to produce equal subjective intensities (10). Performance was quantified through psychometric techniques (Fig. 1B, D). Animals were handled in accordance with standards of the National Institutes of Health and Society for Neuroscience. All protocols were approved by the Institutional Animal Care and Use Committee of the Instituto de Fisiología Celular (UNAM).

### Recordings

Recordings were obtained with quartz-coated platinum-tungsten microelectrodes (2 to 3 MΩ; Thomas Recording) inserted through a recording chamber located over the central sulcus, parallel to the midline. Midbrain DA neurons were identified on the basis of their characteristic regular and low tonic firing rates (1-10 spikes per second) and by their long extracellular spike potential (2.4ms ± 0.4 SD). We furthermore verified that the 22 cells used for the study did show a positive activation to reward delivery in correct (rewarded) trials and with a pause in error (unrewarded) trials. A similar criterion has been adopted in many electrophysiological studies of midbrain DA neurons (28, 29).

### Analysis of behavioral data

Animals performed the task for multiple sessions composed of about 120 trials. Behavioral data were obtained on average from 2226 trials per stimulus class (Fig. 1D). To classify trials according to the RT we defined short-RT trials as trials with RT below the 50th percentile and long-RT trials those with RT above the 50th percentile (Fig. 1E-F).

### Analysis of the firing rate activity

The responses (z-scores) to f_1_ and f_2_ in Fig. 2A-B were standardized with respect to a temporal window preceding the onset of the base stimulus that lasted 500ms and was centered 1000ms after KD. To estimate the temporal profile of the z-score (Fig. 4, Fig. 5A-B left and center, Fig. 5C, and Fig. 6A) we calculated the firing rate for each neuron in 250ms sliding windows shifted every 50ms and standardized it as in Fig. 2A-B. Finally, the responses (z-scores) during the delay period (Fig. 5A-B right) were measured during its entire duration (3s) and standardized as in Fig. 2A-B.

### Latency values

The firing rate time-course of the responses to f_2_ depended on trial uncertainty and trial outcome (Fig. 4). To determine the time of divergence between the two time-courses, we applied a receiver operating characteristic (ROC) curve analysis in each sliding temporal window. This was done within the period lasting from 300ms before f_2_ onset to 200ms after its offset. For each neuron we obtained the normalized firing rate (z-score) in sliding windows of 250ms shifted in 10ms steps. We used the z-scores of all neurons and trials to calculate the ROC curve at each time bin. The area under the ROC curve (AUROC) was used as the index indicating differential neuronal activity across different trial types. Values of the AUROC higher or lower than 0.5 indicated that, at the population level, one type of trial evoked a higher or lower DA response than the other. To determine the statistical significance of the computed AUROCs, we used a permutation test with 1000 resamples. Significance was determined with p<0.05 in 5 consecutive windows and the latency was defined as the first window that met this criterion. A similar analysis was used to determine significant differences in the temporal profile of the normalized activity for trials sorted according to the RT.

### Dependence of the DA activity on f_1_ and class number

In order to search for f_1_-dependent activity we performed two different tests. We first used a linear and a sigmoidal regression analysis (Fig. 6E) to assess whether the activity was monotonic with respect to f_1_. Then, we used a one-way analysis of variance (ANOVA) test to identify any general, non-monotonic relationship between firing rate activity and f_1_. We focused both analyses on the f_1_-stimulation period and WM delay between f_1_ and f_2_. We calculated a mean time-dependent z-score (standardized as in Fig. 2A-B) using a sliding window of 250ms moving in steps of 10ms, from 0.5s before f_1_ onset up to 0.5s after f_2_ offset. Window times with a significant monotonic signal (slope different from zero, p<0.01, for either a linear or sigmoidal fit with Q>0.05) were marked as “significantly monotonic.” Window times where the ANOVA was significant (p<0.05) were marked as “significantly dependent” (11, 19). We then divided the 5s period (from 0.5s before f_1_ onset up to 0.5s after f_2_ offset) into 10 non-overlapping intervals of 500ms and counted the number of windows that were significantly linear in each interval. For each interval, we said that the f_1_ dependence was significantly linear if more than the 40% of windows were significantly linear, and significantly dependent if more than 40% of windows gave a significant ANOVA p-value. Fig. S2A shows the temporal evolution of the p-value resulting from these multiple ANOVA tests. Fig. S2B shows the z-scores in windows in which the dependence was significant. A similar procedure based on the ANOVA test was employed to calculate how the responses depended on the class number. The temporal evolution of the p-value resulting from the ANOVA tests and the z-scores in windows with significant dependence are shown in Fig. S2C-D, respectively.

### Correlations between DA and RT

To obtain the correlations between RTs and DA activity (z-scores) in Fig. 6B-C we employed data from all correct trials independently of the class presented. The z-scores were obtained as in Fig. 2A-B with specific temporal windows for each analyzed event. 250ms after cue presentation (PD), the z-score was obtained using a window of 300ms of width. Similarly, correlations at f_1_ presentation were computed using a window of 480ms of width centered at 240ms after the onset of the stimulus. The z-score used to obtain the correlations during the delay period (Fig. 6C left) was in a window centered 2.5 seconds after the first stimulus offset and of 1 second width. The correlation coefficient between RT and DA was calculated using the MATLAB function “corrcoef”. To verify the significance of the correlations, we performed a permutation test (Fig. 6B-C left) using 10000 randomly shuffled samples. Corrcoef function also provides a p-value for the coefficient. This p-value coincided up to the second decimal point with the one obtained with the permutation test. During the 3s delay period, the temporal evolution of correlations (Fig. 6C right) was studied by obtaining the z-score in 17 windows of width 300ms evenly spaced between 0.5 and 3 seconds after the offset of f_1_. Correlations, which became significant towards the end of the delay, were consistently negative regardless of the number of windows as well as the width of them. Significance was assessed using corrcoef function in MATLAB.

### Bayesian model for the discrimination task

The discrimination task was modeled using a Bayesian framework. The prior probabilities of f_1_ and f_2_ were taken to be uniform and denoted as *P*(f_1_^i^) (i = 1, …, 6) and *P*(f_2_^j^) (j = 1, …, 10), respectively. It was assumed that the animal knew the class structure used in the experiment (Fig. 1B-C), but it had access only to noisy representations (observations) of the two frequencies presented in the trial (denoted by f_1,0_ and f_2,0_). At the onset of the second stimulus, an observation *o*_2_ of the frequency f_2,0_ was obtained from a Gaussian distribution with mean f_2,0_ and standard deviation σ_2_. This noisy information was combined with knowledge of the prior distribution *P*(f_2_^j^) to obtain the belief, or posterior distribution about the value of the second frequency, *b*_2_(f_2_^j^ = *P*(f_2_^j^|*o*_2_) α *P*(*o*_2_| f_2_^j^) *P*(f_2_^j^). The observation of the first frequency f_1,0_, made at the end of the delay period, had to be retrieved from working memory and was indicated by *o*_1_^*^. This observation was also taken from a Gaussian distribution, but with mean f_1,0_ and standard deviation σ_1_. The belief about the value of the first frequency was denoted by *b*_1_(f_1_^i^) = *P*(f_1_^i^ | *o*_1_^*^).

The belief state B(*k*| *o*_1_^*^, *o*_2_) about the class c_k_ = (f_1_^*k*^, f_2_^*k*^) (*k*=1, …, 12; class labels, Fig. 1C) was defined as the set of the posterior probabilities *P*(f_1_ = f_1_^*k*^, f_2_ = f_2_^*k*^ | *o*_1_^*^, *o*_2_) that the class *c*_k_ had been presented in this trial, conditioned to the observations *o*_2_ and *o*_1_^*^. It can be written as:

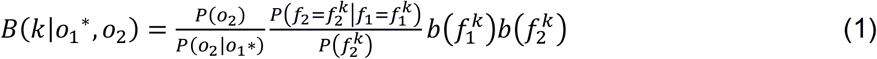

The first factor in the above equation is a normalization factor. The second factor is a transition matrix relating the first and the second stimulation frequencies divided by the prior probability of the second stimulation frequency. Since we assumed that the animal had perfect knowledge of the class structure, the only non-zero matrix elements correspond to the 12 classes *c_k_* of the experiment. Furthermore, since f_2_ can only take two possible values for a given value of f_1_, all non-zero transition probabilities are 0.5. The last two factors then become the beliefs *b*_1_(f_1_^*k*^) and *b*_2_(f_2_^*k*^) about the first and second frequencies being those in class *c*_k_.

The sums of the *B*(*k* | *o*_1_^*^, *o*_2_)’s over classes *k* for each choice, gives the two belief values (f_1_>f_2_ or f_1_<f_2_). These two sums were denoted by *b*(H) (higher choice, f_1_>f_2_) and *b*(L) = 1 - *b*(H) (lower choice, f_1_<f_2_), and choices were made according to the larger of these two, denoted as *b*_c_. The performance can be measured by the fraction of trials of a given class in which the decision is correct.

The two unknown model parameters, the standard deviations σ_1_ and σ_2_, were fitted by minimizing the mean squared error between the model performance and the animal’s performance. We found the optimal parameter values: σ_1_ =5.5Hz and σ_2_ =3.2Hz (see next section for more details about the model fitting procedure).

The statistical uncertainty *U* of a given trial is *U*=1-max[*b*(H), *b*(L)]=1-*b*_c_, which is bounded between 0 and 0.5. The uncertainty of a given class was defined as the average of the value *U* across trials of each class. The uncertainty in hits (or errors) of a given class was defined as the average over the correct (or wrong) trials from each class.

### Bayesian model fitting procedure

Parameter fitting in Fig. 3B and S3 were made using the Simulating Annealing solver in MATLAB. For each RT condition in Fig. S3, 200 fits were performed. The two fits that yielded the lowest error were used to model the monkey’s performance in Fig. 1. Given that 200 adjustments were performed, a distribution for each of the two parameters was available. The mean value for σ_1_ obtained in the short-RT condition was found to be significantly lower than the noise parameter for the long-RT group (one-tailed two sample t-test; p<0.001). The same result was found for the noise parameter σ_2_ (one-tailed two sample t-test; p<0.001).

### Reinforcement Learning Model based on Belief States

To test whether the phasic responses can be attributed to dopamine reward prediction errors (RPEs) we constructed a reinforcement learning model and checked if it was able to reproduce the observed responses. Given that in the task the relevant stimuli are only partially observable (the animal is not aware of the true value of f_1_ and f_2_) we used a belief-state temporal difference (TD) model (similar to that proposed by (5)) to compute reward expectations and simulate the RPEs signaling. This was first implemented in a Bayesian module that works similarly to the Bayesian model described above and uses the same fitted values of the two noise parameters, σ_1_ and σ_2_. This module yields the belief b_1_(f_1_^i^) (i=1, …, 6) about the value of the first frequency, the belief state B(k | o_1_^*^, o_2_) about the class c_k_ = (f_1_^k^, f_2_^k^) (k=1, …, 12) at the time when the second frequency is presented and the belief *b_c_* = [*b*(H), 1-*b*(H)] about which of the two frequencies is higher. Then it transmits these inference results to a TD module that selects actions and generates reward prediction errors (RPEs).

The RL model also uses a fully observable variable *pm* (push movement) that represents the movement towards one of the two buttons. This variable has two possible states: *py* (“push yes”) when the decision is f_1_>f_2_ and *pn* (“push no”) when the decision is f_1_<f_2_. Finally, the reward function, denoted by *r*, is a scalar function that takes two different values for correct and incorrect decisions (see Equation 8).

At the beginning of each trial the TD module calculates the value of the first stimulus as:

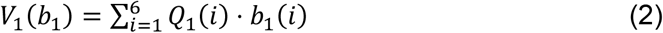

where *Q*_1_(i) is a set of adaptable weights. The RPE at the first stimulus is *δ*(*f*_1_) = *V*_1_(*b*_1_) At the second stimulus the TD module computes the value of the class:

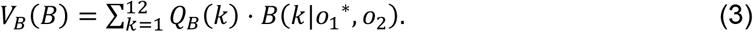

Here the *Q_B_*(k) (k=1, …, 12) are another set of adaptable weights. At the second stimulus the TD module also estimates the value resulting from the comparison of the two frequencies

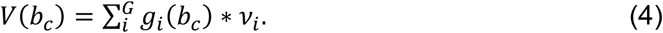

where the *ν_i_* are a set of adaptable weights. The *g_i_* (i=1, …, G) are convenient functions that account for the contribution of the belief *b_c_* to this value. The functions *g_i_* were taken as the following basis functions:

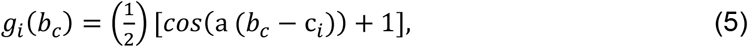

if *c_i_* – 0.1 < *b_c_* < *c_i_* + 0.1 and *g_i_*(*b_c_*) = 0 otherwise. The c_i_’s are G=11 equally spaced centroids in [0, 1] and a = π/0.1.

Given the values *V_B_*(*B*) and *V*(*b_c_*) the RPE at the second stimulus is:

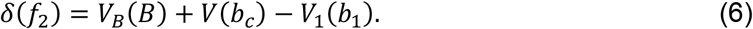

As in the Bayesian model, we assumed that decisions were made according to the larger of the beliefs *b*(H) and *b*(L) about which of the two frequencies was the higher. The value of the response movement when action j is selected is indicated with *V_rm_*(j) to highlight the correspondence between the action selected and the subsequent movement.

The RPE at the response movement is:

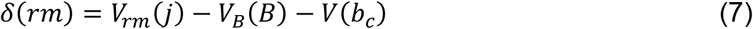

and the RPE at the delivery of reward is:

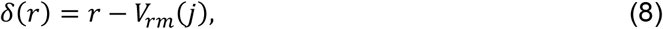

where r =1 for correct discrimination and r = −0.5 otherwise.

The RPEs *δ*(*f*_2_), *δ*(*rm*) and *δ*(*r*) are used to update the adaptable weights *Q_1_*(i), *Q_B_*(k), *ν*_i_, the value of the movement *V_rm_*(j).

We assumed a discount factor *γ* = 1 and used the TD(λ) algorithm to update the weights at the end of each trial. The updating rule is the following:

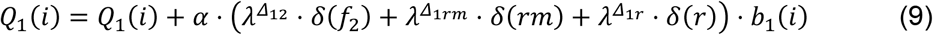

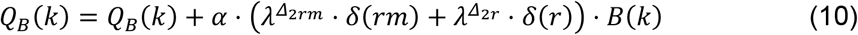

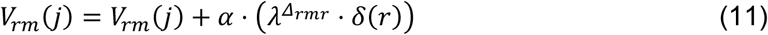

In all the equations above *α* represents the learning rate. The value of *Δ* is the temporal interval between the relevant task events, i.e. the onset of f_1_, the onset of f_2_, the response movement and the reward. To mimic the task temporal structure we took *Δ*_12_ = 30, *Δ*_1*rm*_ = 40, *Δ*_1*r*_ = 45, *Δ*_2*rm*_ = 10, *Δ*_2*r*_ = 15, *Δ_rmr_* = 5 (The temporal intervals are expressed in units of the time step, *dt* = 0.1 s). For the simulation, we use *λ* = 0.95 and *α* = 0.05.

---

**Figure S1.**
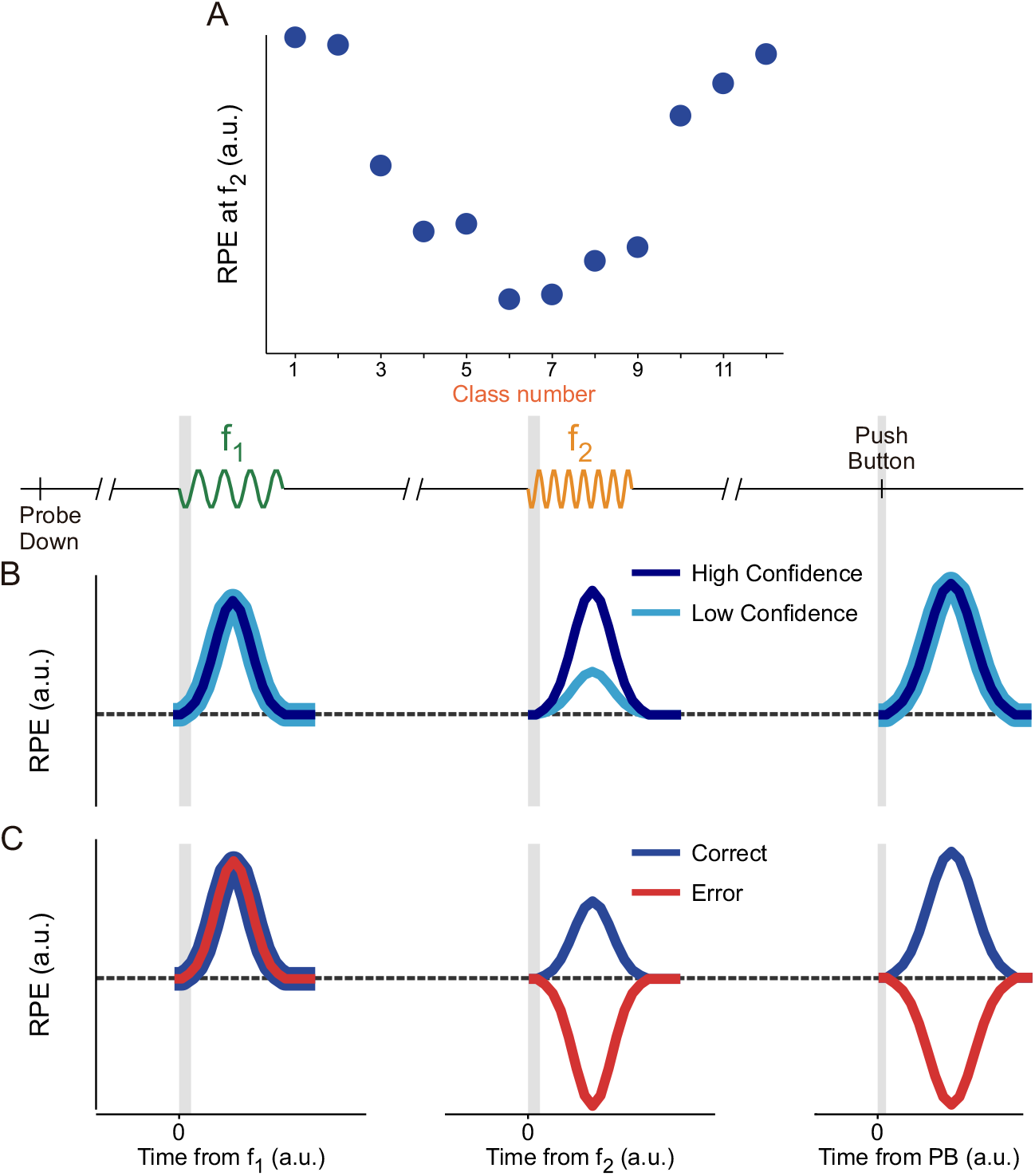
Reward prediction errors predicted by the reinforcement learning model are similar to DA phasic responses (related to Figures 2 and 4). (A) RPE as a function of class number in correct trials, taken at the onset of f_2_. A Gaussian filter was applied to the model predictions. Noise parameters σ_1_ =5.50 Hz for f_1_ sampling and σ_2_=3.2 Hz for f_2_ sampling. (B) RPE signal calculated during 3 different periods with emulated data: f_1_ presentation, f_2_ presentation, and reward delivery after Push Button (PB, decision report). Dark blue curve represents the simulated high confidence group. Light blue curve represents the simulated low confidence group. Left: RPE after the onset of f_1_ does not depend on the confidence level, observable in the overlap between both of our confidence group curves. Center: RPE after the onset of f_2_ depends on choice confidence, favoring the high confidence group. The low confidence group (light blue) reaches less than half the max value observed for the high confidence group. Right: RPE after reward delivery peaks at a value independent of the confidence level since the curves overlap perfectly. (C) RPE calculated during 3 different periods for simulated correct trials (blue curve) and simulated error trials (red curve). Left: RPE after the onset of f_1_ is similar in correct and error trials, since the two curves overlap. Center: RPE after f_2_ onset shows activation in correct trials and a strong depression in error trials (red curve). Right: RPE generated by the model after PB. Correct trials show an increase in RPE, while the error trials show a depression in RPE. The amount of change between error and correct trials is approximately equivalent.

**Figure S2.**
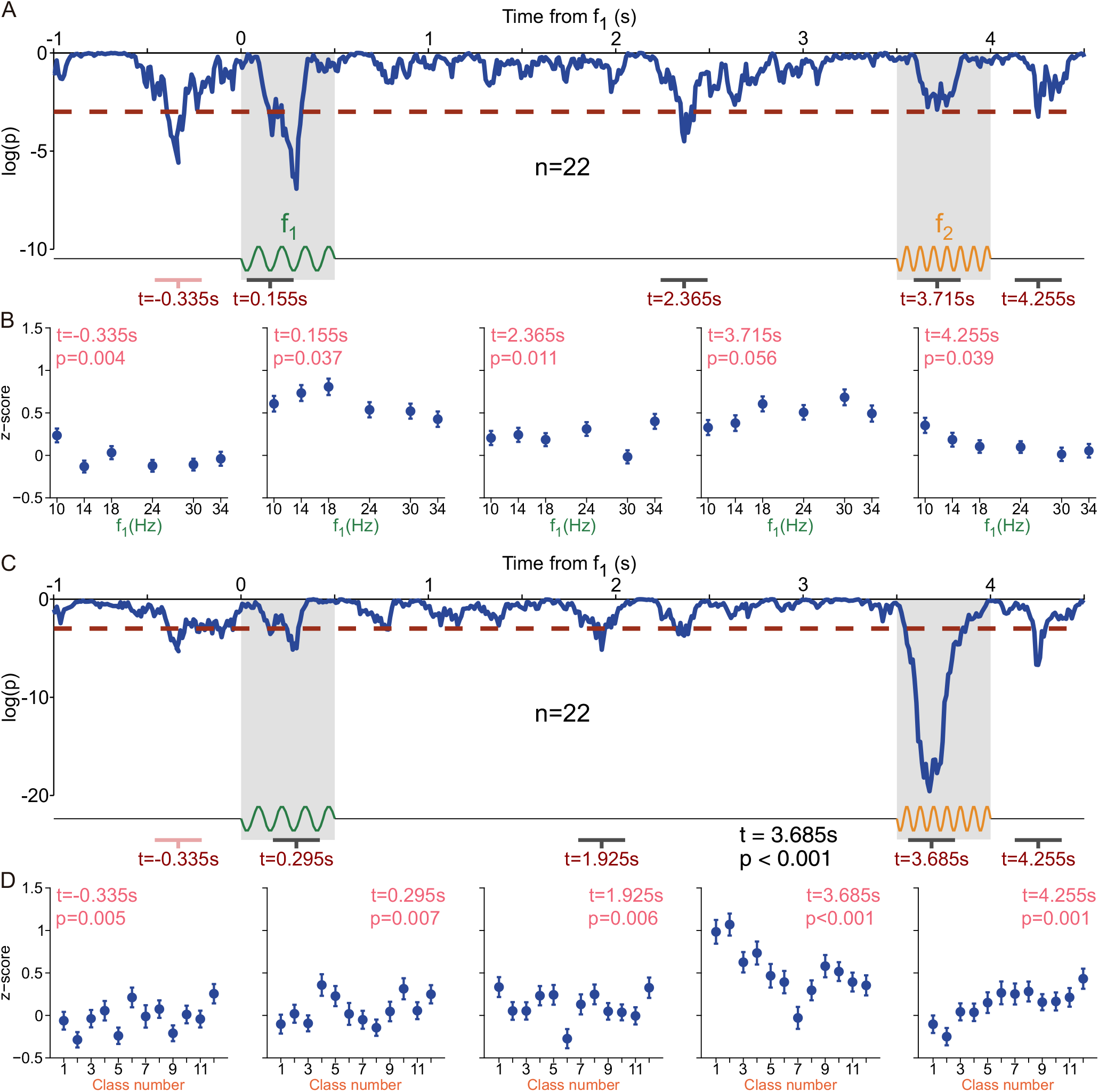
Analysis of the existence of general dependencies on f_1_ and on class number during the trial (related to Fig. 5E). (A) Temporal evolution of the logarithm of p-value (log(p)) resulting from multiple ANOVA tests performed to check the dependence of the z-score on the f_1_ values. We calculated a mean time-dependent z-score (standardized as in Fig. 2A-B) using a sliding window of 250ms shifting in steps of 10ms and performed multiple ANOVA tests sorting the z-score according to the values of f_1_. Values below the red dotted lines are considered as significant (significance is assessed as p<0.05). The p-value intermittently crossed the significance threshold. However, it consistently remained significant only during the presentation of the first stimulus. (B) Average z-score for each f_1_ stimulus (error bars for ±1 SEM) calculated in each of the 5 indicated regions from panel (A). First graphic for the basal period before f_1_ onset, proceeding in increasing chronological order from left to right. Times in the top left corner (pink) and p values (pink) are the results of the ANOVA tests and indicate dependencies on f_1_ when below threshold (red dotted line in panel A). (C) Similar to panel (A) but the p-value was obtained by sorting the z-score according to the class number and running multiple ANOVA tests. (D) Similar to panel (B) but averaging the z-score according to the class number.

**Figure S3.**
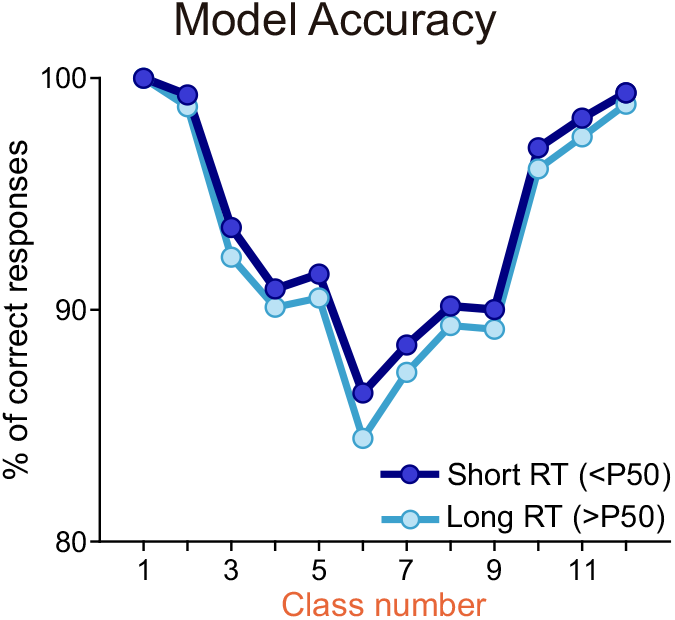
Percentage of correct responses as a function of class number obtained with the Bayesian model for short- and long-RT trials (light and dark blue circles and line, respectively; related to Fig. 1F, left). Model best-fit parameter values were σ_1_=5.28 Hz and σ_2_=3.01 Hz for the short-RT group, and σ_1_=5.385 Hz and σ_2_=3.0 Hz for the long-RT group.

